# Integrated Analysis of HeberFERON-Driven Comparative Proteomic regulation in Glioblastoma Cells U-87MG

**DOI:** 10.64898/2026.04.22.720155

**Authors:** Dania Vázquez-Blomquist, Vladimir Besada, Jamilet Miranda, Yassel Ramos, Caridad Sucel Palomares, Osmany Guirola, Ricardo Bringas, Eva Vonasek, Yolimar Gil, Wendy Pérez, Tamara Díaz, Mauricio Quiñones, Luis Javier González, Iraldo Bello-Rivero

## Abstract

Glioblastoma is a very aggressive brain tumor with few therapeutics’ options. Type I and II Interferons (IFNs) co-formulation HeberFERON has been used in cancer treatment, with promising results in high grade brain tumors. High throughput techniques in easy-to-handle models have been important to interrogate biomolecules changes, describe mechanisms and find pharmacodynamic biomarkers. This study aims to elucidate the effect of HeberFERON over the cell proteome in comparison to its individual IFNs components. Proteomic changes with HeberFERON in the glioblastoma-derived cell line U-87MG, in comparison with individual IFN-α2b and IFN-γ, were studied using a nanoLC instrument EasyLC coupled to Velos Pro mass spectrometer; Maxquant and Perseus were also used. Several enrichment tools, networking analysis and canSAR for drug targets were employed. Translation, RNA processing, mitotic cell cycle, cytoskeleton and chromosome organization, apoptosis, autophagy, DNA repair are enriched to limit cellular growing together with changes in immune response components, supporting HeberFERON as a multitarget treatment. This co-formulation is distinguished at modulating RNA splicing with SMN complex, cytoskeleton organization and microtubule-based movement, nuclear envelope breakdown, DNA conformational changes, and oxidative phosphorylation, with a better drawing of effects over a variety of systems inside the tumoral cell. Together with previous microarray experiment, informative genes and proteins as pharmacodynamic biomarkers for antiproliferative effects showed up (ex. STAT1/2, CENPE, ATRIP, MAP1B, LIMA1, VCP, several ribosomal, spliceosome and proteasomal complexes proteins). This study complements transcriptomic and phosphoproteomic previous experiments in this model and underscore HeberFERON as a glioblastoma therapeutic.

## 1 Introduction

Glioblastoma (GBM), are invasive and aggressive tumors of the brain with a median survival of 12-14 months [1]. First line of treatments, with resection by surgery followed by radiotherapy and chemotherapy, have not significantly increased overall patients’ survival. As a new therapeutic approach, we are proposing the use of a co-formulation of recombinant Interferons (IFNs) type I [alpha (α)2b] and type II [gamma (γ)], named HeberFERON [2]. Interferons have demonstrated effects over cancer cells using direct and indirect mechanisms [3]. Their crosstalk in pathways and the membrane association of their receptor chains, molecularly support the use of combinations [4]. HeberFERON has shown antiproliferative effects in different glioma cell lines, including the widely used U-87MG [5] and patient-derived clones [6]. Moreover, nude mice models with xenografted U-87MG-based tumors also show the decrease of tumoral volume and increase of survival in animals receiving HeberFERON or temozolomide in comparison to a placebo group (*unpublished results*). This cellular model has been very useful to understand the mechanisms at different levels beneath this treatment. Previous gene expression profile study by a microarray was carried out, also comparing with individual IFNs treatments [5] giving a view of compared regulation at transcriptomic levels. A phosphoproteomic study was recently described [7] as another layer of regulation. In all the cases co-formulation was used at the IC50 for 72h. In the present study, we aimed to elucidate the effect of HeberFERON over the cell proteome in comparison to its individual IFNs components.

## 2 Materials and Methods

### 2.1 Reagents

Recombinant interferons, rIFN-α2b and rIFN-γ, and the pharmaceutical co-formulation (HeberFERON) were produced at CIGB, Havana, Cuba.

### 2.2 Cell lines and culture conditions

Human glioblastoma cell line U-87MG (ECACC 89081402, Salisbury, Wiltshire, UK) [8] was maintained in complete medium MEM (Sigma, US), supplemented with 10% Fetal Bovine Serum, 2mM glutamine and 50 µg/ mL of gentamicin (All Gibco, US) in a humidified atmosphere of 5% CO_2_ at 37°C. Cells were grown at a cellular density of 35 000 cells/ cm^2^.

### 2.3 Experiment Design for Proteomic

An experiment with three replicates per U-87MG cells treatment condition was designed. Groups included: untreated cells (cc1-cc3), rIFNα2b treatment (a1-a3), rIFNγ treatment (g1-g3) and treatment with the combination HeberFERON (ag1-ag3). After 24h, complete MEM containing recombinants rIFNα2b, rIFNγ or HeberFERON at IU/mL equivalent to IC_50_ of HeberFERON were added to the experimental groups. Cells were incubated for another 72h; untreated cells received only complete MEM. 10 million cells per sample were washed once on-flask with cold PBS and scrapped. Three washes with cold PBS and centrifugations at 100 g for 10min at 4 °C completed the procedure until obtaining a protein pellet.

### 2.4 Sample preparation and MS analysis

Protein extracts were prepared in 500 µL of 6 M guanidine hydrochloride, 50 mM DTT, 100 mM HEPES at pH 8.5, containing protease and phosphatase inhibitors. After centrifugation at 60000 g the supernatant was kept for one hour at 37 °C and cysteines were later blocked with 25 mM acrylamide. Protein digests (70 μg) were obtained with Lys-endopeptidase (WAKO, Japan) and trypsin (Promega, USA) at 1/100 and 1/50 enzyme: substrate ratio, respectively. Desalted samples were analyzed with a nanoLC instrument EasyLC (Thermo, USA) coupled to a Velos Pro mass spectrometer (Thermo, USA). The chromatographic separation was performed through a column Easy 75 µm x 100 mm (Thermo, USA) in a three steps gradient of acetonitrile/ 0.1% formic acid 2-20% in three minutes, 20-40% in 77 minutes and 40-80% in 23 minutes at 300 nL/min. Mass spectra were acquired from 350 up to 1650 Th, at 1900 V capillary voltage, and resolution of 60000 in the Orbitrap analyzer. HCD (higher-energy collision dissociation) MS/MS spectra were obtained for the first 19 more intense signals. An isolation window of 2 m/z, 35V activation energy, 50 ms activation time, 7500 resolution of the Orbitrap was selected. Automatic gain control (AGC) was set to 1×10^6^ (MS) and 5×10^4^ (MS/MS).

### 2.5 Protein identification and quantification

MS/MS database searching was performed with Maxquant (version v1.6.0.16) on Swissprot database. Four peptide modifications were included: methionine oxidation, deamidation of asparagine and glutamine and protein N-terminal acetylation as variable modifications, and propionamide cysteine as fixed modification. Mass tolerance of 10 ppm for the precursor ion and 4.5 ppm for the fragments was considered. Target decoy method and FDR below 5% were used for protein identification.

For free labeling quantification program Perseus v1.6.0.7 was used. We considered non-less than three replicates per protein. We performed a paired test for each condition (IFNα, IFNγ and HeberFERON) in comparison to the control condition, rejecting the null hypothesis for those proteins with a change factor ≥ 1.5 and a p-value < 0.05. FDR of 5% was also considered. Differentially expressed proteins were reported as log2 (IFNα/CC; IFNγ/CC; HeberFERON/CC). This program was also used to find proteins related with different biological processes of interest.

### 2.6 Gene Ontology Analysis and Functional classification

Database for Annotation, Visualization and Integrated Discovery (DAVID 6.8, https://david.ncifcrf.gov/home.jsp) [9] was used for Gene Ontology, Functional annotation and Clustering analysis taking the identified proteins as a background of analysis. ToppFun (from ToppGene Suite https://toppgene.cchmc.org/) [10], and Metascape (https://metascape.org/gp/index.html#/main/step1) [11] were also used as analysis tools (until May, 2025) to identify molecular functions, biological processes and pathways altered in the proteome of U-87MG cells due to the effect of treatments (p< 0.05). Venn diagrams were constructed to know common and different proteins among treatments using the online resource from VIB /UGent Bioinformatics & Evolutionary Genomics (Gent, Belgium; http://bioinformatics.psb.ugent.be/webtools/Venn/). A list of differentially expressed proteins (DEPs, Supplementary Table 1S), was used for gene ontology analysis.

### 2.7 Network analysis

Cytoscape (v3.2.1) [12] with BisoGenet (v3.0.0) [13] as plugin were used to constructed and visualize networks; MCODE (v1.4.1) plugin [14] to find highly interconnected proteins on network.

### 2.8 Analysis by canSAR

Bioinformatics tool canSAR [15] allowed to know the studied drug targets contained in DEPs changed by HeberFERON.

### 2.9 Effects of HeberFERON combined with protease inhibitors over U-87MG cell line

HeberFERON was assayed in nine 1:2 dilutions (from 100000 to 312 IU/mL) in an antiproliferative assay set up. Sulphorhodamine B [16] method was performed in 96 flat bottom plates seed with 10^5^ U-87MG cells/mL in 100 µL, for 72h. Two proteases’ inhibitors were tested. Bestatin is a Leucin aminopeptidase (LAP) inhibitor, including LAP3; it is a dipeptide analogue with a Ki of 9nM and 3μM for LAP and APN proteases, respectively. E-64 is a potent irreversible inhibitor against general cysteine proteases, including Cathepsin S. Bestatin was assayed using a two-fold dilution curve from 300 to 1.1 μM and E-64 from 800 to 12.5 μM [17–20]. We performed three independent experiments with two interplate replicates for each dilution. Program Calcusyn (Calcusyn BioSoft Version 2.0, UK) was used for IC_50_ calculation. This program draws a regression line, plotting log [concentration] versus log [Fa/FU] (FU: relation of absorbance at 492 nm of a given dilution respect to the absorbance of the cell control after medium values subtraction; Fa=1-FU) with an r>0.90. Combinations of HeberFERON with Bestatin and E-64 were performed using a range of doses as follows: HeberFERON (0-1250-2500-5000-12500-25000 IU/mL) and Bestatin (0-7.8-15.6-31.25-62.5-125 µM); HeberFERON (0-6250-12500-25000-50000-100000IU/mL) and E-64 (62.5-125-250-500-1000µM). Graphs showing the cellular affected fraction at growing HeberFERON concentrations, fixing Bestatin or E-64 concentrations, were constructed. We also calculated the Dose-Reduction Index (DRI) values for synergistic drug combinations at a relation 1:40 (Bestatin: HeberFERON) and 1:100 (E-64: HeberFERON). The DRI is a measure of how much the dose of each drug in a synergistic combination may be reduced at a given effect level compared with the doses for each drug alone [21].

## 3 Results

### 3.1 Differentially expressed proteins in response to IFNα, IFNγ and HeberFERON

LC-MS/MS proteomic analysis identified a total of 2207 proteins, 756 (34 %) were significantly DEPs with respect to untreated cells: 230 with IFNα, 213 with IFNγ and 514 with HeberFERON (Fig. 1A). Venn diagram shows common and different proteins among the three treatments. Twenty-nine are common among the three treatments. Although DEPs are shared between groups, 362 proteins are differentially expressed only by HeberFERON. Principal Component Analysis (PCA) showed replicates per group clustered closer and well differentiated from the other groups of treatments replicates (Fig. 1B).

**Fig. 1.**
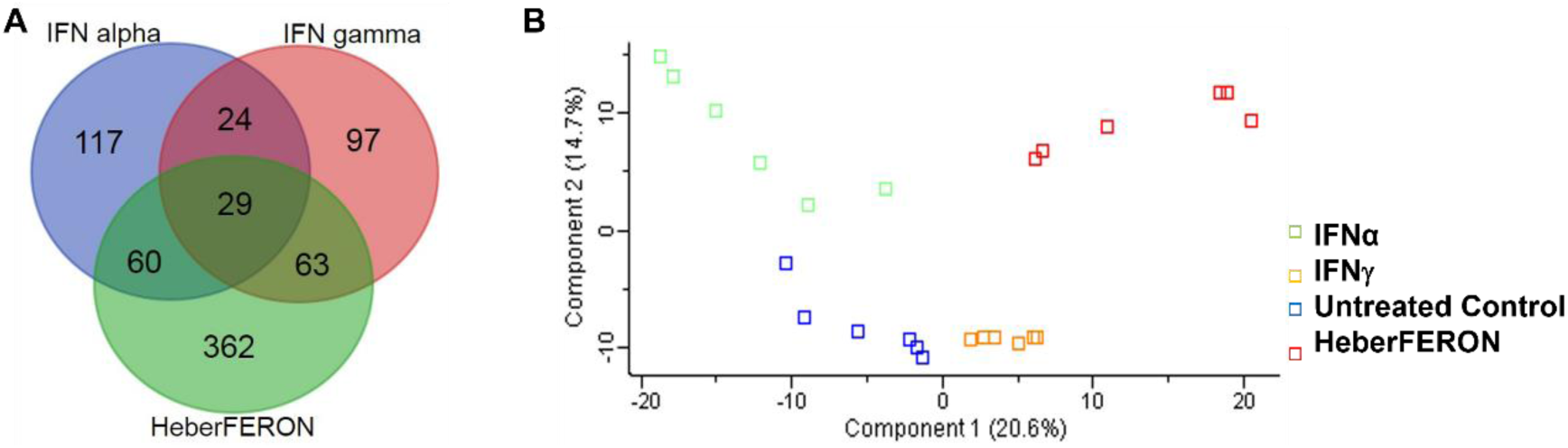
A) Venn diagram among the DEPs in groups treated with IFN alpha, IFN gamma and HeberFERON. B) PCA analysis of proteome data in all the replicates of the experiment groups treated with IFNα, IFNγ and HeberFERON and the Untreated control.

### 3.2 Biological functions of DEPs by HeberFERON

The analysis with the 514 DEPs by HeberFERON using DAVID software shows several enriched terms from pathways, processes and molecular functions related to Interferon Signaling, Canonical glycolysis, Cytokine Signaling in Immune system, Innate immune response and Response to virus, Mitotic Metaphase and Anaphase (with additional related terms such as: Separation of Sister Chromatids and Mitotic G2-G2/M phases), Proteasomal protein catabolic process and Proteasome complex, PKR-mediated signaling, Apoptosis, Microtubule and Microtubule-based process, Antigen processing-Cross presentation and Spliceosomal complex assembly (Fig. 2A). A parallel enrichment analysis by Toppfun gives similar results for proteins regulated by HeberFERON. Fig. 2B shows biological functions more represented in the data using Gene Ontology information. In addition to the biological processes mentioned using DAVID, proteins changing their abundance with HeberFERON participate in translation/ribosome biogenesis, rRNA metabolic process, chromosome organization, cytoskeleton organization, microtubule cytoskeleton organization involved in mitosis, spindle organization, protein folding, mitochondrial ATP synthesis coupled electron transport and DNA repair.

**Fig. 2.**
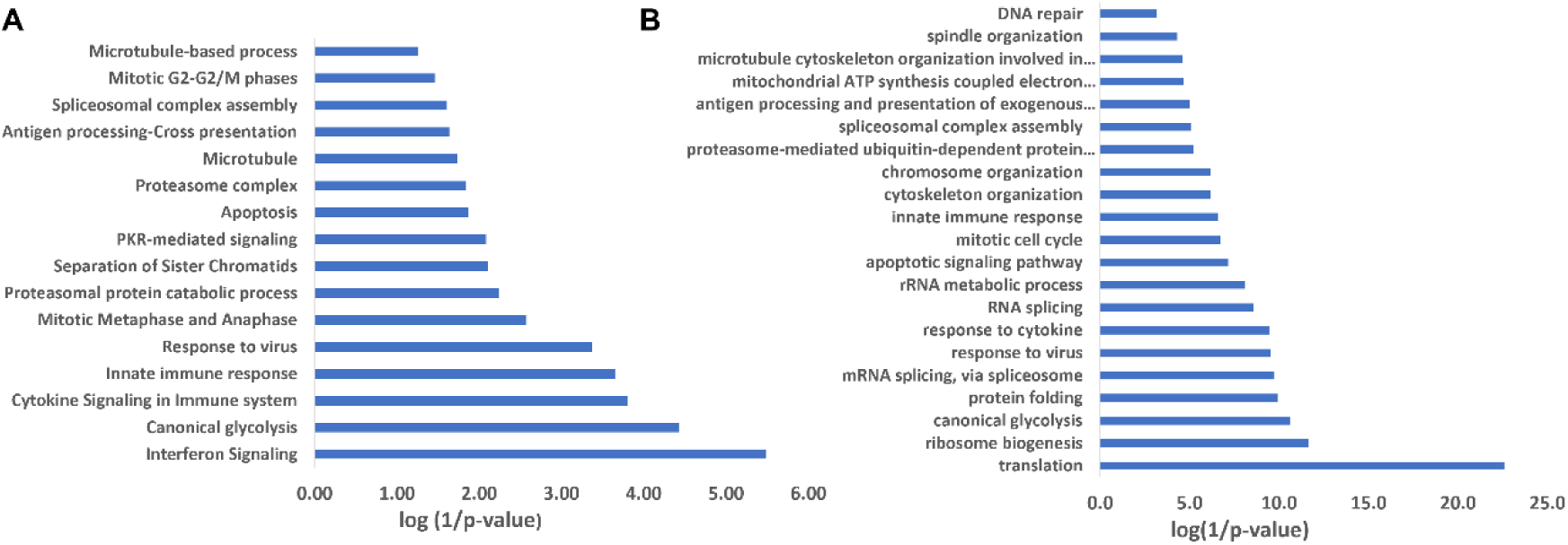
Enriched biological functions from proteins differentially expressed by HeberFERON, IFNα2b and IFNγ. A: Analysis by DAVID using DAVID terms *vs* (log(1/p-value) for HeberFERON; B: Analysis by Toppfun using Toppfun terms vs log(1/p-value) for HeberFERON.

### 3.3 Biological functions of DEPs by HeberFERON in comparison to individual interferons

HeberFERON regulated the same 89 and 92 proteins as IFNα and IFNγ, respectively, but 362 proteins were only regulated by the co-formulation. Beyond individual proteins, the inspection of the Top 100 most represented biological processes in the list of DEPs, showed there are some that are shared by the three treatments but with a higher enrichment after HeberFERON treatment such as: Cytokine Signaling in Immune system, Interferon Signaling, Metabolism of RNA (rRNA processing, mRNA Splicing), Spliceosome, Translation, Apoptosis (Regulation of Apoptosis), Ribosome, Mitotic Cell Cycle (Mitotic Metaphase and Anaphase, Separation of Sister Chromatids, G2/M Transition, Cell Cycle Checkpoints, APC/C-mediated degradation of cell cycle proteins), Antigen processing-Cross presentation (Antigen processing: Ubiquitination & Proteasome degradation), Antiviral mechanism by IFN-stimulated genes and response to virus (PKR-mediated signaling, ISG15 antiviral mechanism),Proteasome (Proteasome degradation), purine and pyridine ribonucleotide metabolic processes, Carbon metabolism, generation of precursor metabolites and energy, Autophagy (Macroautophagy, Selective autophagy) and other (Fig. 3 and Supplementary Table 2S). But there are also processes that HeberFERON shares only with IFNα or with IFNγ. As examples, protein folding and refolding, ribosomal large subunit biogenesis, folding, assembly and peptide loading of class I MHC, nucleosome assembly and Mitotic prophase are shared by HeberFERON and IFNγ. Otherwise, with IFNα, HeberFERON shares the mRNA processing, NADH regeneration/canonical glycolysis/glucose catabolic process to pyruvate, Apoptotic execution phase, and antiviral innate immune response. The most interesting thing is that there are GO terms that only appeared with HeberFERON treatment like SMN complex, regulation of RNA splicing and mRNA splicing via spliceosome, Initiation of Nuclear Envelope (NE) Reformation and Nuclear Envelope Breakdown, regulation of cytoskeleton organization and microtubule polymerization or depolymerization, microtubule-based movement, regulation of chromosome organization, DNA conformation change, DNA repair, positive regulation of cysteine-type endopeptidase activity, for example.

**Fig. 3.**
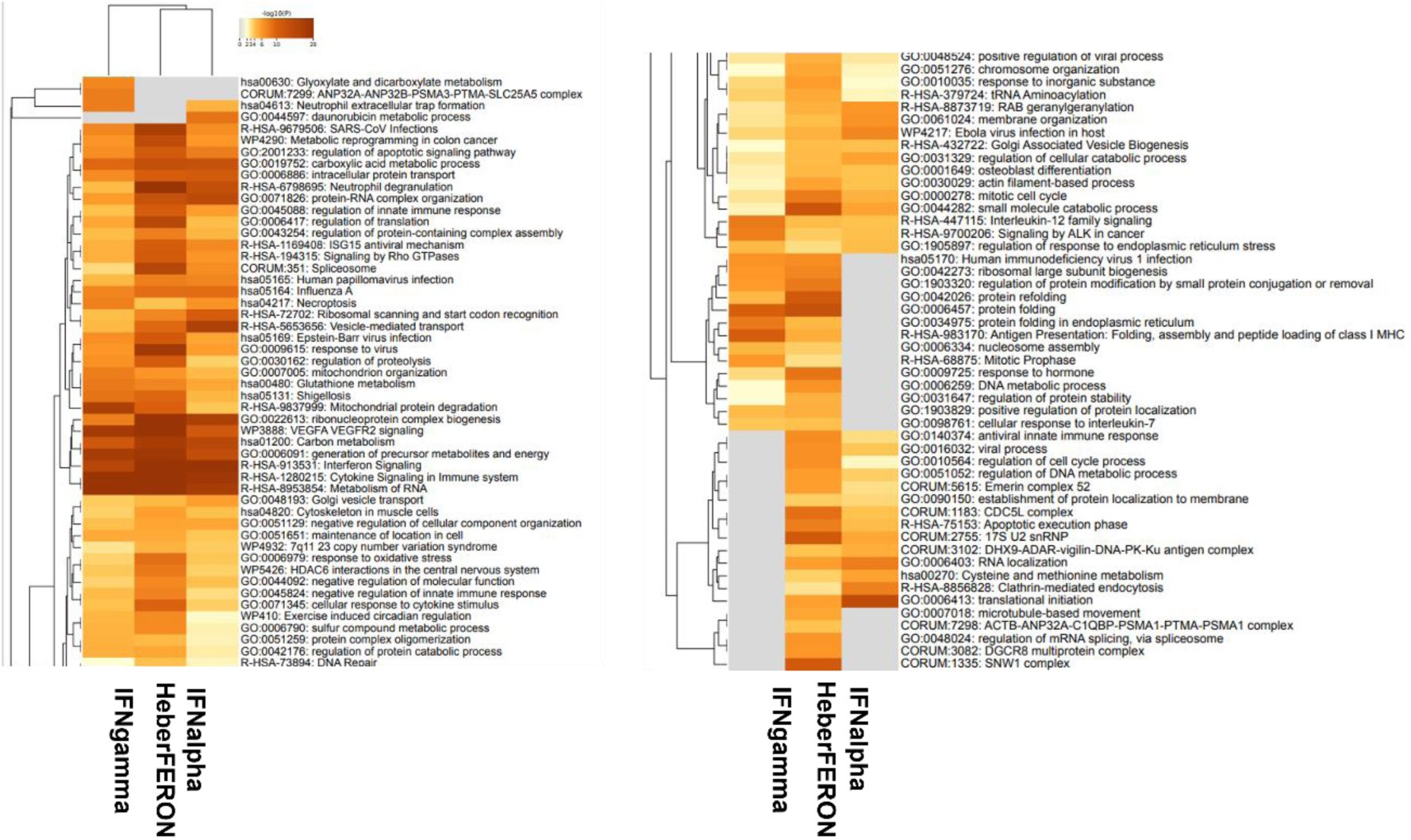
Top enriched ontology clusters across U87-MG cell line treated with IFNalpha, IFNgamma and HeberFERON for 72h using Metascape. The most 100 enriched biological processes (-log10(P)) in U87-MG after IFNs treatment are shown.

We selected some biological processes to visualize with Volcano plots using Perseus. It is evident the increase of DEPs participating as “Interferon related” proteins and in “Ribosome”, “Proteasome”, “RNA processing” or “Cell Cycle” after HeberFERON treatment (Fig. 4). HeberFERON showed synergic behavior increasing the levels of some “classical” IFNγ- and IFNα-stimulated proteins such as HLA-I and II proteins, GBP1 and GBP2 or ISG15, respectively. It also increased the very important IFN receptors cascades signal transducers STAT1 and STAT2, but not STAT3 (Supplementary Fig. 1A). STAT1 expression increased more than 4 times by HeberFERON and STAT2 is up regulated 2 times respect to basal levels only by HeberFERON. As for STAT2, MX1 and OAS3 were found upregulated only by HeberFERON.

**Fig. 4.**
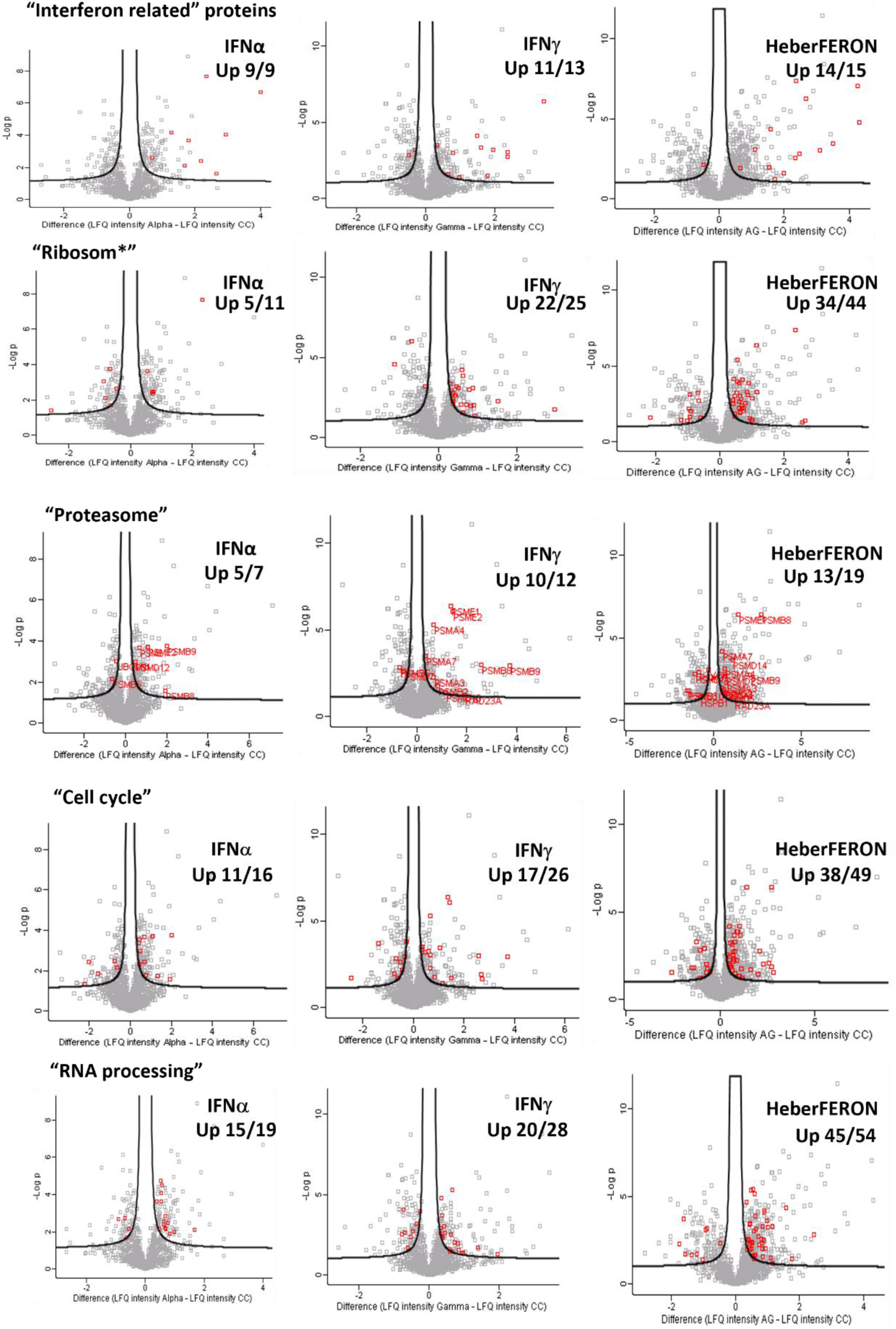
Volcano plots in Perseus. DEPs by each Interferon treatment were investigated in biological categories such as “Interferon related” proteins, “Ribosom*”, “Proteasome”, “RNA processing” and “Cell Cycle”. The upper-right corners of graphs show the number of up-regulated proteins out of the number of total proteins related with the searching term.

From the 49 proteins regulated in “Cell Cycle” category, 17 proteins are from Proteasome. Components of the Immunoproteasome [PSMB8, PSMB9 and the PA28 activator complex gene encoding subunits PSME1/E2] are regulated by HeberFERON as by IFNγ and IFNα. PSMA3, -A4, - A7 and PSMD7 are components regulated by IFNγ & HeberFERON in similar ways. PSMA1, PSMB3, PSMC2, PSMD2, PSMD3, PSMD5, PSMD10, PSMD11 and PSMD14 are only regulated by HeberFERON (Supplementary Fig. 1B).

A similar behavior was found for ribosomal proteins from both Ribosome subunits, where RPL6, RPL7, RPL9, RPL13 and 13A, RPL15, RPL22, RPL23 and 23A, RPL26, RPL27 and 27A, RPL29, RPL34, RPL37A, RPS3, RPS9, RPS10, RPS15A, RPS27 and 27L are only regulated by HeberFERON. Moreover, ribosomal proteins RPL10, RPL12 and RPS11 were regulated by HeberFERON in a different direction as by IFNγ or IFNα (Supplementary Fig. 1C).

### 3.4 Interconnection among DEPs

The interconnection among proteins regulated by IFNα, IFNγ and HeberFERON is high as only 4.8 % were disconnected from the principal network (36/756) (Supplementary Fig. 2), where proteins only regulated by HeberFERON represents about 50 % in the overall data (yellow nodes in the network). Highly interconnected clusters representing biological processes related to Translation/Ribosome, Proteasome, and Spliceosome were identified with 36.1, 16.4, and 5.8 MCode scores (Fig. 5A).

**Fig. 5.**
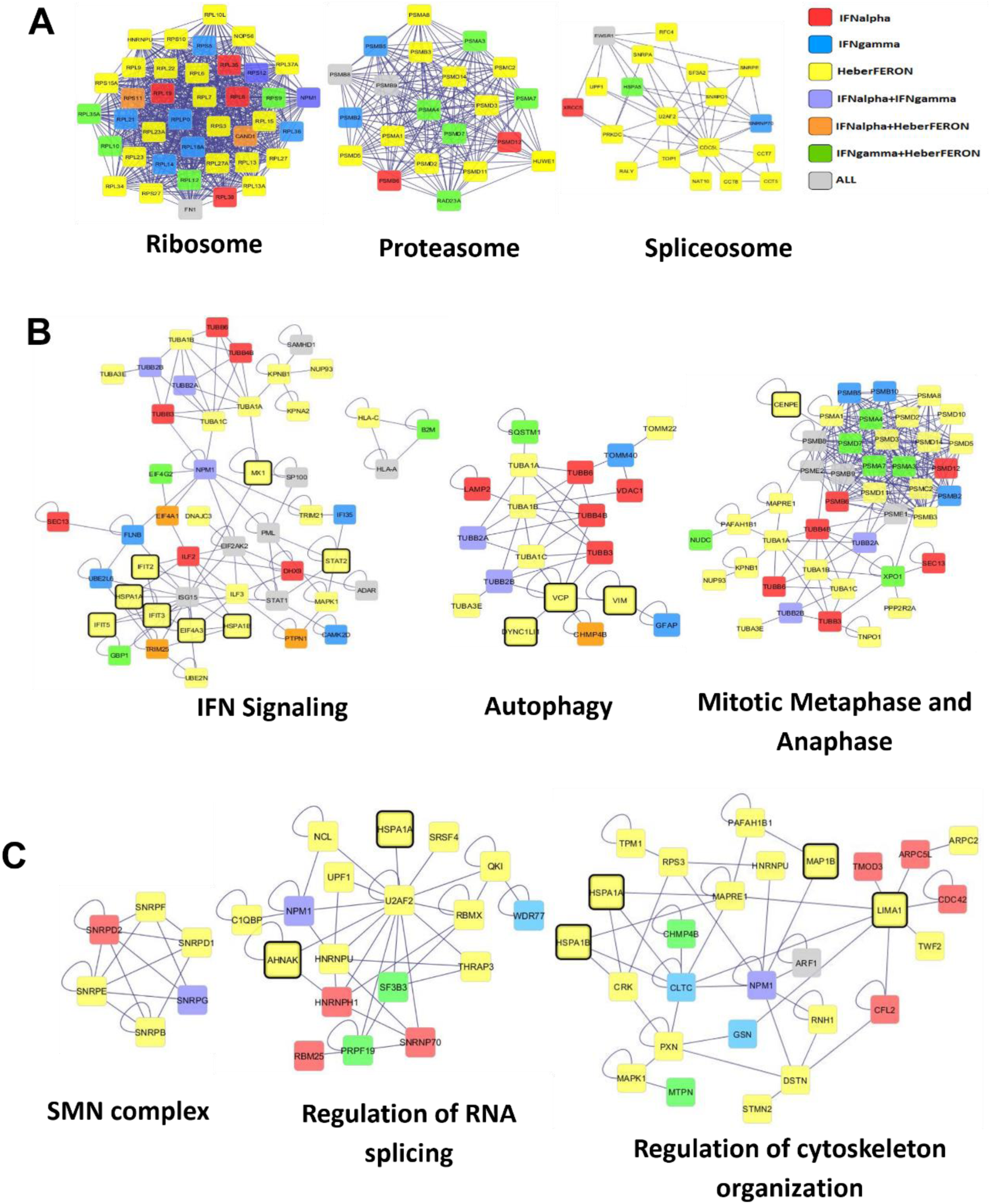
interconnected Clusters from networks formed with all DEPs in this proteomic experiment. A) The most interconnected clusters related to Ribosome, Proteasome, and Spliceosome using MCode plugin; B) Interconnection of DEPs from enriched biological processes for the three interferons but with higher enrichment for HeberFERON; C) Interconnection of DEPs from enriched biological processes only for HeberFERON. Some biomarkers with relevance in previous experiments are indicated. Legend indicates treatments by color.

Additionally, we constructed networks with components of biological processes enriched for the three treatments but higher for HeberFERON, including Interferon Signaling, Mitotic Metaphase and Anaphase and Autophagy (Fig. 5B). It is important to note that important ISG like STAT2, MX1, EIF4A3 and IFIT2/3/5 are only regulated by HeberFERON. Other proteins participating in Autophagy (VCP, VIM, DYNC1LI1) and Mitotic Metaphase and Anaphase (CENPE) have been previously described for this co-formulation in microarray and phosphoproteomic experiments. In relation to Mitotic Cell Cycle and cell cycle checkpoints we also found interesting biomarkers like BANF1, ATRIP and GORASP2. The last two are only down-regulated by HeberFERON. BANF1 and NUP93 also participated in Nuclear Envelope Breakdown, a process that it is only enriched by the co-formulation. Fig. 5C represents the interconnections among DEPs participating in selected processes only regulated by HeberFERON such as SMN complex, regulation of RNA splicing, and regulation of cytoskeleton organization. In these networks DEPs previously found in microarray and phosphoproteomic experiments appeared, like AHNAK participating in the Regulation of RNA splicing and HSPA1A/B, LIMA1, MAP1B in cytoskeleton organization.

### 3.5 Convergence between Microarray and Proteomic experiments for HeberFERON

Taking the data from a microarray experiment with identical design we looked at the common genes and proteins regulated by HeberFERON in Microarray & Proteomic experiments. The interception between both data was 338 gene/proteins. The selection of some biomarkers such as the STAT1&2, and other involved in Antiviral response, Antigen Presentation, Proteasome, Translation, Spliceosome, Cell Cycle, Cytoskeleton Organization, Chromosome Organization and Autophagy, showed its relevance on the effect of HeberFERON in relation to individual IFNs which could be relevant as potential pharmacodynamic biomarkers (Fig. 6).

**Fig. 6.**
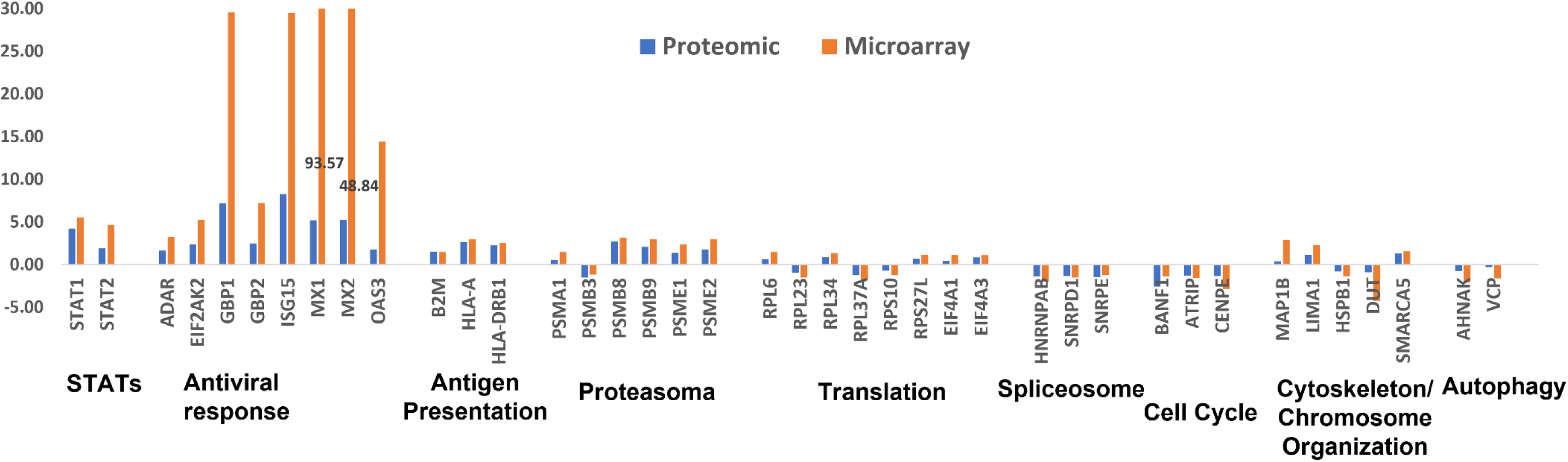
Gene and protein expression levels in U-87MG samples treated with HeberFERON. Fold change of genes and proteins in HeberFERON-treated cultures in relation to control cells from comparative microarray and proteomics experiments is shown. Biomarkers are organized by process of interest.

### 3.6 canSAR Analysis

To find appropriate targets to be used for the treatment of glioblastoma together with HeberFERON we performed an analysis by canSAR obtaining a list of screened active compounds targeting proteins regulated by HeberFERON in this experiment. Table 1 shows selected targets organized by Protein Families. Several protein family members are represented in the list; some of them are already decreased by HeberFERON including tubulin family members, ATRIP and VCP. It was interesting to note the large representation of members of peptidase family, including proteasome components, cathepsins and LAP3; most of them increased expression after HeberFERON treatment.

**Table 1.**
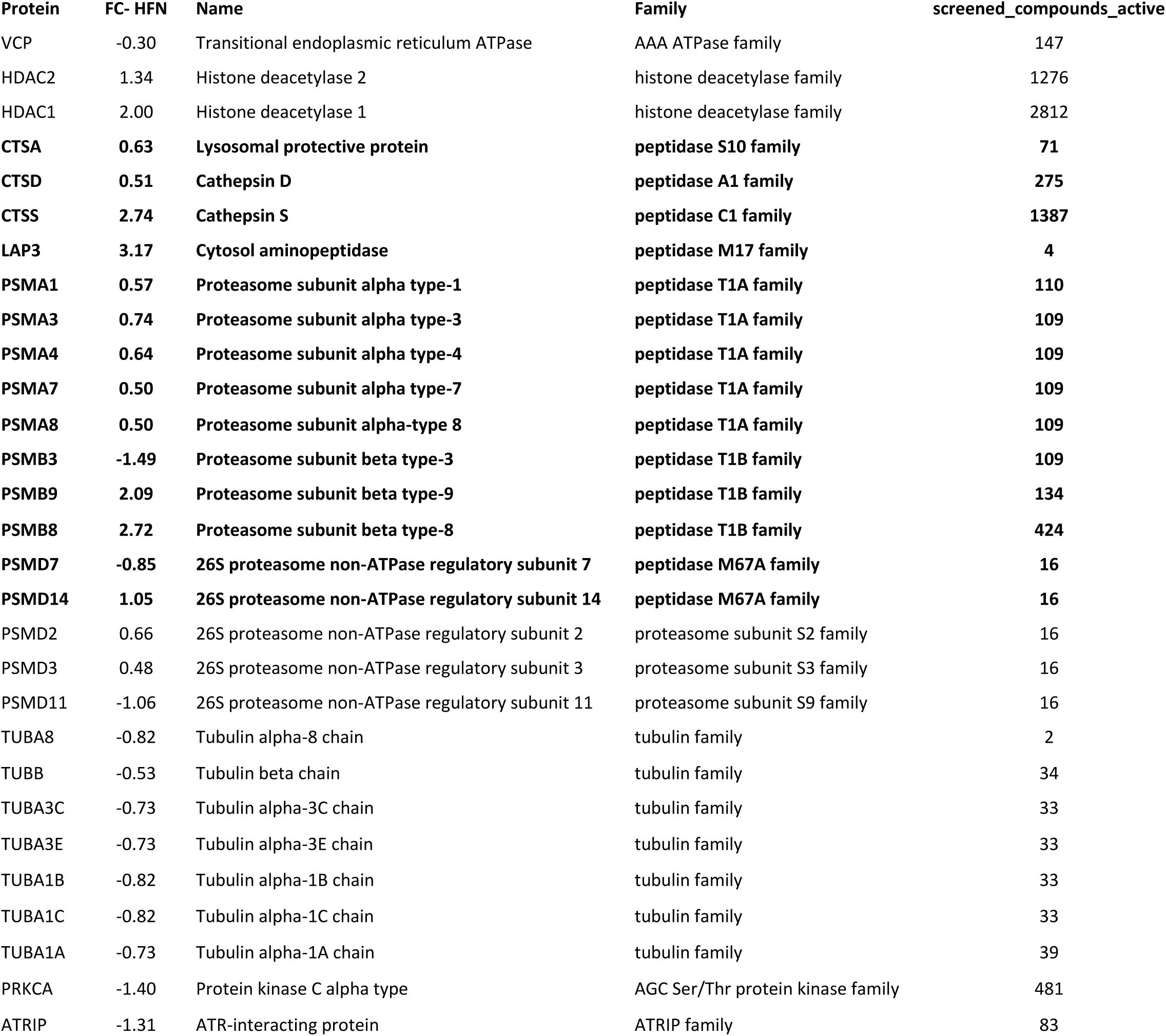
Exploration of the number of screened compounds against proteins regulated by HeberFERON using canSAR. A selection of proteins organized by protein Family is shown. The Table also shows the name (official gene name), fold change with HeberFERON (FC_HFN), protein and protein family names and the number of screened active compounds (screened_compound_active). Proteins from peptidases families are in Bold.

| Protein | FC- HFN | Name | Family | screened_compounds_active |
| --- | --- | --- | --- | --- |
| VCP | -0.30 | Transitional endoplasmic reticulum ATPase | AAA ATPase family | 147 |
| HDAC2 | 1.34 | Histone deacetylase 2 | histone deacetylase family | 1276 |
| HDAC1 | 2.00 | Histone deacetylase 1 | histone deacetylase family | 2812 |
| **CTSA** | **0.63** | **Lysosomal protective protein** | **peptidase S10 family** | **71** |
| **CTSD** | **0.51** | **Cathepsin D** | **peptidase A1 family** | **275** |
| **CTSS** | **2.74** | **Cathepsin S** | **peptidase C1 family** | **1387** |
| **LAP3** | **3.17** | **Cytosol aminopeptidase** | **peptidase M17 family** | **4** |
| **PSMA1** | **0.57** | **Proteasome subunit alpha type-1** | **peptidase T1A family** | **110** |
| **PSMA3** | **0.74** | **Proteasome subunit alpha type-3** | **peptidase T1A family** | **109** |
| **PSMA4** | **0.64** | **Proteasome subunit alpha type-4** | **peptidase T1A family** | **109** |
| **PSMA7** | **0.50** | **Proteasome subunit alpha type-7** | **peptidase T1A family** | **109** |
| **PSMA8** | **0.50** | **Proteasome subunit alpha-type 8** | **peptidase T1A family** | **109** |
| **PSMB3** | **-1.49** | **Proteasome subunit beta type-3** | **peptidase T1B family** | **109** |
| **PSMB9** | **2.09** | **Proteasome subunit beta type-9** | **peptidase T1B family** | **134** |
| **PSMB8** | **2.72** | **Proteasome subunit beta type-8** | **peptidase T1B family** | **424** |
| **PSMD7** | **-0.85** | **26S proteasome non-ATPase regulatory subunit 7** | **peptidase M67A family** | **16** |
| **PSMD14** | **1.05** | **26S proteasome non-ATPase regulatory subunit 14** | **peptidase M67A family** | **16** |
| PSMD2 | 0.66 | 26S proteasome non-ATPase regulatory subunit 2 | proteasome subunit S2 family | 16 |
| PSMD3 | 0.48 | 26S proteasome non-ATPase regulatory subunit 3 | proteasome subunit S3 family | 16 |
| PSMD11 | -1.06 | 26S proteasome non-ATPase regulatory subunit 11 | proteasome subunit S9 family | 16 |
| TUBA8 | -0.82 | Tubulin alpha-8 chain | tubulin family | 2 |
| TUBB | -0.53 | Tubulin beta chain | tubulin family | 34 |
| TUBA3C | -0.73 | Tubulin alpha-3C chain | tubulin family | 33 |
| TUBA3E | -0.73 | Tubulin alpha-3E chain | tubulin family | 33 |
| TUBA1B | -0.82 | Tubulin alpha-1B chain | tubulin family | 33 |
| TUBA1C | -0.82 | Tubulin alpha-1C chain | tubulin family | 33 |
| TUBA1A | -0.73 | Tubulin alpha-1A chain | tubulin family | 39 |
| PRKCA | -1.40 | Protein kinase C alpha type | AGC Ser/Thr protein kinase family | 481 |
| ATRIP | -1.31 | ATR-interacting protein | ATRIP family | 83 |

### 3.7 Effect of combination of HeberFERON with peptidase inhibitors

As a proof of combining HeberFERON with inhibitors targeting proteins that increased their expression with HeberFERON, we selected two model peptidases, cathepsin S and LAP3, and their inhibitors E-64 and Bestatin. The improved effect combining HeberFERON with Bestatin was observed (Fig. 7A) when fixing a concentration of Bestatin and plotting the affected fraction (Fa) in the evaluated range of doses for HeberFERON. For all the doses of Bestatin there was an improved antiproliferative effect of the combination with HeberFERON respect to HeberFERON alone at the same dose. The highest effects are observed in the range of concentration of HeberFERON from 1250 to 5000 IU/mL; for some Bestatin doses (125, 62.5, 15.6 µM) the effect is even slightly higher at a concentration of HeberFERON of 12500 IU/mL. Calcusyn program also calculate de DRI for each drug for Fa from 0.02 to 0.99. These DRI derives a new calculation for IC50 for the combination taking into account the Bestatin, meaning that to account a 50 % of cell growth inhibitions we can reduce an original dose of Bestatin from 182.4 µM to 62.6 µM.

**Fig. 7.**
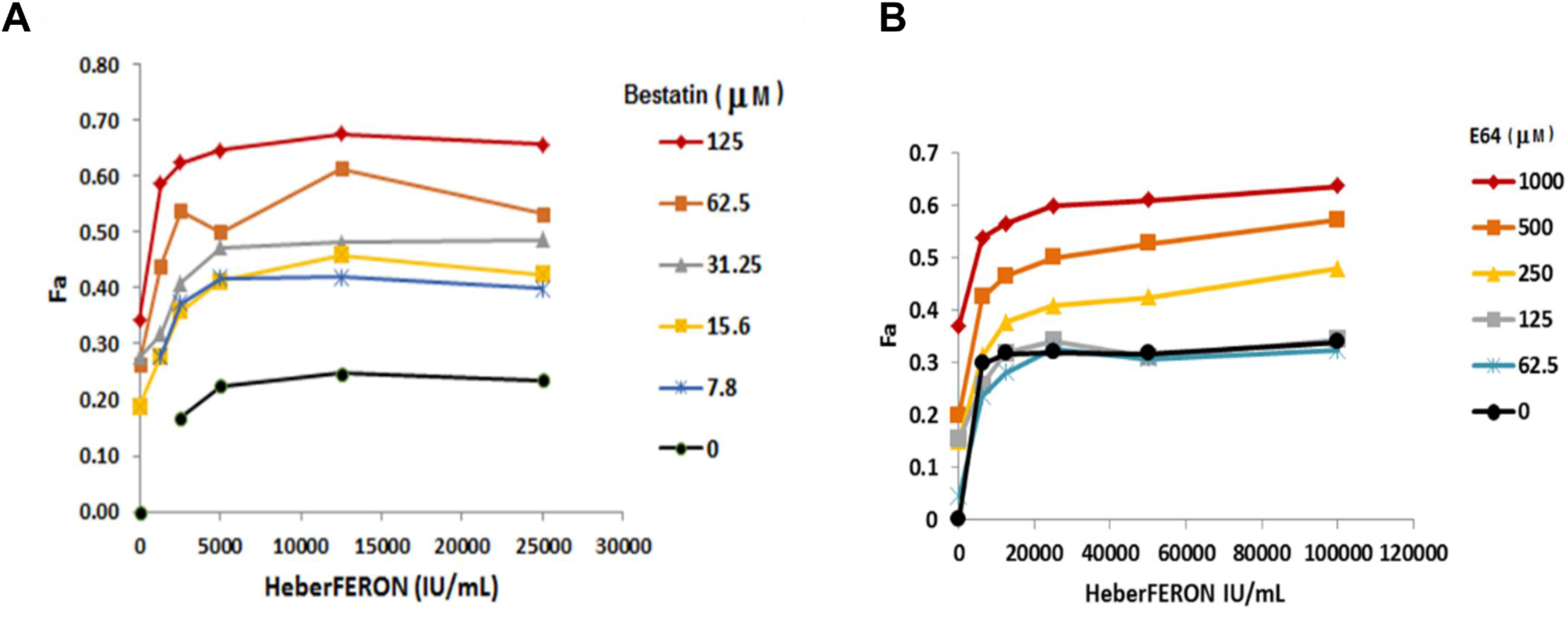
Fraction affected (Fa) of cells in function of HeberFERON concentrations (IU/mL) for fixed Bestatin (A) and E-64 (B) doses (in µM).

The improved effect at combining HeberFERON with E-64 was observed (Fig. 7B) when fixing a concentration of E-64 and plotting the Fa in the evaluated range of doses for HeberFERON. From 0.25mM of E-64 there was an improved antiproliferative effect for the combination with HeberFERON respect to HeberFERON alone at the same dose. The growth inhibition of 50 % of cells is only achieved at the highest doses of E-64 (0.5 and 1mM) and increased with addition of higher doses of HeberFERON. The calculation for IC50 for the combination, taking into account the E-64 showed a reduction of 4.7 times of the doses when combining with HeberFERON. It means that to account a 50 % of cell growth inhibitions we can reduce an original dose of E-64 from 1987.7 µM to 421.2 µM.

## 4 Discussion

HeberFERON, as a co-formulation of synergic proportions of IFNα2b and IFNγ [2], is expected to show synergistic and additive effects, taking into account the crosstalk between both receptors and signaling pathways [4]. As we previously published in a comparative gene expression microarray experiment with identical design [5], at the protein level there also are shared proteins between individual IFNs treatments and HeberFERON, 29 proteins shared by the three, and in a way that point to HeberFERON as a “new interferon”, 362 proteins only regulated by the co-formulation.

Glioblastoma is a very deadly brain tumor with treatment schedules without a deep impact on the overall survival [22]. New therapeutics should cross blood-brain barriers and then deal with the high tumor heterogenicity, glioblastoma stemness and the immunosuppressive tumoral microenvironment [1]. In this sense, HeberFERON shows promising possibilities because the pleiotropic effects of interferons and their impact through direct and indirect immune system-supported mechanisms.

HeberFERON regulated more than 500 proteins from several biological processes and protein complexes like Response to viruses, Immune response, Translation and Ribosome, RNA processing and Spliceosome, Proteasome, Mitotic Cell Cycle, cytoskeleton and Microtubule-based process, chromosome organization, Apoptosis, Autophagy, and DNA repair. This experiment also showed the contribution of canonical glycolysis and mitochondrial ATP synthesis coupled electron transport as processes that could also limit the cellular growing. Oxidative phosphorylation is a process down-regulated only by HeberFERON. Synergic effects are evident in Volcano plots contrasting “Interferon related” proteins and proteins from Ribosome, Proteasome, RNA processing or Cell Cycle regulated by IFNα, IFNγ and HeberFERON.

The interconnection among proteins is high in networks and highly connected clusters of DEPs related to IFN signaling, Ribosome, Spliceosome, Proteasome, cytoskeleton organization, autophagy and Cell Cycle appeared. It is not completely a surprise, HeberFERON could have an effect over those processes as a combination but, undoubtedly, this comparative proteomic experiment complements the former transcripts-based exploration to draw the main processes and players in its effect over U-87MG. A subtractive hybridization experiment using HEp-2 cells treated with HeberFERON also showed regulated genes participating in immune system and antigen presentation (B2M, HLA-C, HLA-B, HSP90AB1, HSPA5 and HSPD1), in Cytoskeleton regulation (ACTB, ACTG1, RHOA) and a high proportion in protein translation and ribosomal biogenesis (RPL10A, RPL24, RPL4, RPL7, RPS16, RPS19, RPS21, RPS27A, RPS3A, EEF1A1, EIF4A3, rRNAs 18S and 28S) [23]. These processes seem like universal mechanisms for HeberFERON effects over tumoral cell lines.

DEPs linked to antiviral effects or antigen processing and presentation on an MHC context for an effective immune response are presented in the IFN-related response, and regulated in higher magnitudes by HeberFERON. IFNs activate an antiviral host defense pattern in humans fibrosarcoma cell line HT1080, hepatoma cells Huh7, lung carcinoma cells A549 and other cell types [24–26]. Proteins like PKR, OAS proteins producing 2’5’-oligoadenylates, the guanosine triphosphatases Mx, the IFN-induced guanylate-binding proteins (GBP) and ISG15 are among those components of the innate immunity, also used as IFN-pharmacodynamic biomarkers [27].

This link could be the explanation for the effect of HeberFERON over basal cell skin carcinomas during COVID-19 pandemia intervention [28]. Then the immune response should play an essential role as part of the indirect mechanism to control tumor growing and spreading as it does for viral control [29]. Interferons are emerging as an attractive therapeutic in Cancer because it can act over different actors in the tumoral microenvironment and contribute to alert immune system by increasing innate immunity, the membrane expression of tumor associated antigens and their antigen processing and presentation or other immune related regulatory biomarkers [29]. Further preclinical and clinical translational investigations would be necessary to validate these hypotheses.

In the canonical JAK-STAT type I &II interferons signaling pathways, STATs are key elements [30]. IFNα induces the phosphorylation of both STAT1 and STAT2, producing STAT1 homodimers and STAT1:STAT2 heterodimers (complexed with IRF9 form ISGF3 that recognizes IFN-stimulated response elements (ISRE)). IFNγ induces the phosphorylation of STAT1 and promotes the formation of STAT1 homodimers, which recognize γ-activated sequences (GAS)[30]. We previously showed a balance favoring the increase of STAT1 gene instead STAT3 gene in U-87MG treated with the co-formulation HeberPAG [31]. Here, we also obtained an increase in protein STAT1 with all IFNs, and no changes of STAT3 supporting our previous findings [31]. STAT1 is believed to play an important role in growth arrest and apoptosis, and to act as a tumor suppressor[30]. Hartman *et al* demonstrated that the DNA-binding behavior of STAT1 homodimers is not conserved between the different IFN treatment conditions. This introduces another mechanism for the regulation of STAT1–DNA interactions and target gene selection [32]. These authors also found that several of the genes proximal to the STAT1 and STAT2 were involved in the regulation of proliferation and apoptosis [32]. Outstandingly, STAT2 is only regulated in a positive direction by HeberFERON. The meaning of this finding could be related to the antiproliferative response or to the known effect of STAT2 in the IFN signaling control through USP18 [33].

Translation and RNA processing, as well as, Ribosome and Spliceosome as complexes appeared as highly enriched in HeberFERON treated samples. Synchronized synthesis of Ribosomal proteins (RP) and rRNA in G1 prepare cells for mitosis; raw material for ribosome biogenesis is expressed in a coordinated fashion. Proteins in large (RPL6, L7, L9, L13, L13A, L15, L22, L23, L23A, L26, L27, L27A, L29, L34, L37A) and small (RPS3, S9, S15A, S27, S27L) ribosome subunits are only regulated by HeberFERON. Moreover, the type of regulation for proteins RPL10, RPL12 and RPS11 is opposite with HeberFERON and with IFNγ and IFNα, respectively. This property of the combination was previously reported in HEp-2 cell line [23]. Mammalian ribosomes consist of four RNA species and 79 ribosomal proteins (RPs) where most RP genes are coordinately expressed at the mRNA level. Nucleolar assembly begins at the early G1 phase of the cell cycle and it is a hub of ribosomal DNA transcription and rRNA biosynthesis. The newly formed rRNAs together with RPs constitute the building block of the ribosomal machinery. It is also reported that formation of RNP complexes can influence cell growth and apoptosis [34]. HeberFERON could interfere with ribosomal biogenesis, rRNA processing and translation by an unbalanced expression of RP and rRNA quantities or because RPs also participate in extra-ribosomal processes, as it was already suggested in HEp-2 model [23]. In the 362 DEPs only regulated by HeberFERON, SMN complex, regulation of RNA splicing and positive regulation of translation are biological processes down-regulated only by the co-formulation. SMN complex participates in the biogenesis of spliceosome small nuclear ribonucleoproteins, functioning in the assembly, metabolism, and transport of several ribonucleoproteins [35].

Moreover, the participation of Proteasome complex in the mechanism is also evident contrasting Microarray and Proteomic experiments. This complex participates in the antigen processing and presentation on MHC context, in the degradation of components of cell cycle while mitosis proceeds and in the degradation of unfolded proteins, as part of mechanism for the resolution of cellular stress. Components of the Immunoproteasome [PSMB8, PSMB9] and the PA28 activator complex gene encoding subunits [PSME1/E2] are regulated by HeberFERON as by IFNγ and IFNα. IFNs regulate the ubiquitin-proteasomal system at different levels, improving the immune responsiveness of target cells; immunoproteasome and/or proteasome activator PA28 are required for the generation of certain viral epitopes [36]. The 26S proteasome is a highly ordered multicatalytic protein complex composed of the 20S core and 19S regulatory complexes [37]. The 20S core has four rings of 28 non-identical subunits; two rings composed of seven alpha subunits (PSMA1-A7) and two rings with seven beta subunits (PSMB1-B7). PSMA1 increased its expression only by HeberFERON and PSMA3, PSMA4, PSMA7 by the co-formulation and IFNγ; the expression of the PSMB3 subunit is only decreased by HeberFERON. In the other hand, the 19S regulatory complex is composed of a core containing six ATPase subunits and two non-ATPase subunits, and a cap containing up to 10 non-ATPase subunits. Proteasomes cleave peptides in an ATP/ubiquitin-dependent process in a non-lysosomal pathway. In this complex, there are several subunits that change their levels only by HeberFERON. Of the 19S Base, the ATPase PSMC2 (Rpt1) and the non-ATPase PSMD2 (Rpn1) increase their expression. In an initial step of the assembly of a Base subcomplex, an intermediate module composed of PSMD10:PSMC4: PSMC5:PAAF1 is formed, which probably assembles with another module composed of PSMD5:PSMC2: PSMC1:PSMD2. The chaperones in these complexes, PSMD10 and PSMD5, decrease and increase their expression, respectively. From the 19S “lid” there is an increase in the non-ATPases PSMD3 (Rpn3) and PSMD14 (Rpn11) and a decrease in PSMD11 (Rpn6) and PSMD7 (Rpn8); PSMD3, PSMD14 and PSMD7 are part of the association between the “lid” and the Base and additionally, HSP90 and RPN6 (PSMD11) participate in the regulation of the association between the 20S and 19S complexes. The expression of all the subunits of the Proteasome is very well coordinated and regulated to produce the correct amount of each one. Proteasome assembly is a highly energetic process, then it can compromise cellular survival and in this sense proteasome inhibitors have been studied in glioblastoma although mutational and epigenetic makeup can affect their effects [38]. He et al identified a long noncoding RNA (malate dehydrogenase degradation helper /MDHDH) which is downregulated in glioblastoma. The mechanism of this lncRNA is to bind to MDH2 (malate dehydrogenase 2) and PSMA1, accelerating the degradation of MDH2 what changes the mitochondrial membrane potential and NAD+/NADH ratio, obstructing glycolysis in glioma cells [39]. HeberFERON increased both PSMA1 and MDH2. PSMB3 was identified, together with CHCHD4/SPDYE5/HSPA1, with high expression in cancer and with a significant positive correlation with respect to glioblastoma growth and patient survival *in vivo* [40]. HeberFERON decreased PSMB3 and HSPA1. In a systemic analysis of PSMD family members, Li et al showed levels of PSMD 10 and 11 were higher in GBM than in normal brain tissues [41] and the interferon co-formulation decreased the expression of both proteins. Whether the effect of individual components or changes in the stoichiometry and regulation of proteasome proteins contribute to HeberFERON effects is also interesting for a deeper study.

Microarray experiment found HeberFERON-DEG participating in the mitosis (mainly in Prometaphase) but it also regulated different players until anaphase. Proteomic can explain the processing of mitotic proteins by the Proteasome complex for phase transitions. In fact, volcano plot showed 49 proteins regulated in Cell Cycle category and 17 proteins are from Proteasome. Among the higher enriched biological processes for HeberFERON, we found Mitotic Metaphase and Anaphase, Cell Cycle Mitotic, Separation of Sister Chromatids, G2/M Transition and Cell Cycle Checkpoints also with a high contribution of proteasome proteins, together with ATRIP, BANF, CENPE, DYNC1LI1, DYNLL2, GORASP2, KPNB1, NUP93, and components of cytoskeleton including tubulin complexes. The first three biomarkers are found to be downregulated at gene and protein levels.

CENPE protein, participating in spindle checkpoint, is downregulated but other CENP-family proteins or those regulated by FOXM1-PLK1 axis does not appear regulated. The importance of cell cycle in the effect of HeberFERON was corroborated because several components are regulated by changes in phosphorylation of sites in CENPA and INCENP, for example [7]. Moreover, FACS experiments showed time and dose dependence of cell cycle arrest after HeberFERON treatment what is evident as early as 24h. Then, we could think the decreased expression in cell cycle-related genes at 72h is due to cell cycle arrest at earlier times, contributing to diminish the number of cells. But after treatment for 72h with HeberFERON, proteomic showed other players related to Mitosis. Due to inherent delays in translation, protein maturation, and differential degradation rates, proteomic profiles may reflect transcriptional activity from earlier time windows rather than concurrent mRNA. Different kinetic and regulations at gene and protein levels could account for these differences [42]. BANF1 function is related to nuclear assembly during mitosis [43]. Among the processes only down-regulated by HeberFERON we found Nuclear Envelope Breakdown that occur during mitosis to nuclear membrane and pore complexes disassembling [44]. Not only BANF1 was downregulated but also PRKCA associated with Akt-mTORC1 activation. Moreover, in tumor tissues it is significantly associated with poor survival and negatively correlate with immune cell infiltration [45]. Knockout of these genes activates antitumor immune responses mediated by cGAS-STING pathway, resulting in an immune-activating tumor microenvironment including increased CD8+ T cell infiltration and the decrease of myeloid-derived suppressor cell enrichment. IFNγ and HeberFERON decrease its expression. ATRIP is a DNA damage checkpoint protein present in the centrosomes of cells undergoing mitosis [46]. The DNA damage checkpoint halts the cell cycle in response to DNA damage until the injury is repaired. ATRIP expression is known to be essential for the vitality of undamaged, proliferating, or damaged cells. As for the former is also downregulated only by HeberFERON. GORASP2 function is related to establish the structure of the Golgi apparatus [47]. During mitosis, the Golgi undergoes extensive fragmentation through a multi-step process that allows the proper partitioning and inheritance to daughter cells. Golgi fragmentation is required not only for inheritance but also for mitotic entry itself, as its blockage results in cell cycle arrest in G2 [48]. This protein is also downregulated only by HeberFERON.

As part of cytoskeleton that participate in the chromosome segregation mitosis, we found two components of the cytoplasmic dynein complex, DYNC1LI1 and DYNLL2, regulated by HeberFERON. Regulation of cytoskeleton organization, microtubule-based movement, and regulation of microtubule polymerization or depolymerization are biological processes downregulated only by HeberFERON displaying components like CENPE, MAP1B and LIMA1. These last two DEPs, also upregulated at gene expression and/or with several phosphosites modified after HeberFERON treatment [5, 7]. Their functions and possible contributions have been discussed elsewhere [7].

As in phosphoproteomic experiment, it is also evident HeberFERON regulates several rescues signaling by autophagy and DNA repair. Proteins like VCP, VIM, DYNC1LI1, SQSTM1 and RHEB, appeared regulated in Autophagy/macroautophagy/selective autophagy processes. The first DEPs were discussed before [7] and RHEB participates through the mTORC1 signaling pathway. For DNA Repair, biomarkers like NPM1, PRKDC, RAD23A, UPF1, RFC4, SMARCA5, UFL1, NUCKS1 and ATRIP appeared regulated; some of them were discussed before [7].

This comparative proteomic study supports pleiotropic HeberFERON effects, showing biological processes and signaling contributing the most to the effect of this combination of interferons in U-87MG. The role of cell cycle and cytoskeleton organization is strengthened but many other processes contributing to synthesize molecular components for new cells are put in more value for interferon direct effects over cellular growth. The immune system as a protagonist is again showed then surely having impact over the tumors and their microenvironments. Although not all the biomarkers into biological processes are the same, there are some that looks as informative pharmacodynamic genes and proteins for antiproliferative effects (ex. STAT1/2, CENPE, ATRIP, MAP1B, LIMA1, VCP and several ribosomal, spliceosome and proteasomal complexes proteins). AHNAK protein deserve a closer study in the future as a biomarker regulated at different levels, mostly changing the phosphorylation of many phosphosites [7].

This study has technical and biological limitations. Using Orbitrap Velos Pro technology we were only able to detect 2207 that limit the overall interpretation of this data. While this study provides a comprehensive overview of proteomic changes induced by HeberFERON in comparison with individual IFNs, and there are convergences in processes and biomarkers with previous studies at gene and phosphoproteomic levels, functional validation of individual proteins is not part of this publication and should be addressed in future mechanistic studies.

Although these limitations, proteomic experiments are not only useful for mechanistically understanding of HeberFERON effects and pharmacodynamic biomarkers proposals but also to find new targets to hit. canSAR analysis pointed to several proteins that have been object of drug therapies; some of them are already down-regulated by HeberFERON (ex. Tubulin family members, PSMB3, PSMD11, PRKCA and ATRIP). It was interested to note there are several peptidases that are up-regulated. Some of them participate in antigen processing and cross-presentation, apoptosis, aminoacidic metabolism, cellular redox control (CTSD, CTSS, LAP3) and proposed as targets in Cancer [49–51]. We assayed the combination of aminopeptidase inhibitor and HeberFERON as a prove of concept. The lysosomal cysteine proteinase Cathepsin S could be a potential target for anti-cancer therapy, including glioblastoma [50]. Its inhibition by Z-FL-COCHO (ZFL) or LHVS in U-251MG and U-87MG glioblastoma cell lines induced autophagy and subsequent apoptosis through PI3K/AKT/mTOR/p70S6K and JNK signaling pathways [49]. Here we used the general cysteine protease inhibitor E-64 [52] and although the IC50 in U-87MG cells is high and no more than a 65% of inhibition is achieved, a higher impact on cellular growth is observed with increasing quantities of HeberFERON using from 0.5 to 1mM of E-64; the combination of E-64 and HeberFERON accounted for a reduction in 4.7 times of E-64 dose to achieve the IC50. Otherwise, leucine aminopeptidase 3 (LAP3) has been target of new inhibitors as anti-cancer therapy and there is a high expression in glioblastoma [51]. Bestatin is a dipeptide specific for this protease and a general leucine aminopeptidase inhibitor [53]. Similar to E-64, there is an increase in the growth inhibitory effects at combining HeberFERON with Bestatin in the dose ranges assayed and a reduction of 2.9 times of the Bestatin dose to achieve the IC50 in these cells. Based on these preliminary experiments HeberFERON can be also used combined with other inhibitors as it is shown for LAP3 and cathepsin S inhibitors.

Although this comparative proteomic experiment was only carried out in the model cell line U-87MG, the results support HeberFERON as an interesting multitargeted therapeutic alternative to treat glioblastoma because the activation of direct effects over cancer cells but also indirect mechanisms involving the immune system and the tumoral microenvironment. This study permitted to propose several pharmacodynamic biomarkers for HeberFERON; targets that could be hit in combination with HeberFERON were also pointed as possible based in the proof of concept with proteases inhibitors. Future translational experiments should be conducted; however, our findings represent part of the HeberFERON preface in preclinical models.

## 5 Data Availability

All data of proteomic results generated during this study are included in this published article and supplementary information files. Any additional requested information would be available from the corresponding author upon reasonable request.

## 6 Credit authorship contribution statement

Conceptualization, D.V.-B., V.B., L.J.G. and I.B.-R.; Data curation V.B., C.S.P; Formal analysis, D.V.-B., V.B., J.M., R.B., C.S.P., O.G.; Funding acquisition, V.B. and L.J.G.; Investigation, D.V.-B., V.B., J.M., R.B., I.B.R; Methodology, D.V.-B., V.B., Y.R., E.V., Y.G., M.Q., W.P., T.D.; Project administration, L.J.G. and I.B.-R.; Writing-original draft preparation, D.V.-B., V.B.; Writing/review and editing, L.J.G. and I.B.-R. All authors have read and agreed to the published version of the manuscript.

## 7 Funding

This work was supported by the Center for Genetic Engineering and Biotechnology, Havana, Cuba and IVIC Institute in Caracas, Venezuela.

## 8 Declaration of competing interest

The authors declare no conflict of interest in publishing these data. These data or part of have not been published in any other scientific journal before.

## Supporting information

Supplementary Tables for the manuscript

## Acknowledgments

The authors are grateful to the institutions where the experiments were carried out. To DrC. Isel Pascual in the Center for Proteins studies at the Faculty of Biology in Havana University for providing the Bestatin and the E-64.

## Supplementary Figures and Tables Captions

**Table 1S.** Results of the comparative proteomic experiment. Majority protein IDs in Uniprot, Protein names, Gene names (Official gene symbols), Difference respect to cell control, -Log Student’s T-test p-value. (Excel document extra file, Sheet 1)

**Table 2S.** Top enriched GO biological processes (logP_) across proteomic results by treatments with IFN alpha, IFN gamma and HeberFERON using Metascape. Data is organized to show those shared processes among treatments and the ones only enriched with HeberFERON at the bottom. (Excel document extra file, Sheet 2)

**Supplementary Fig. 1S.**
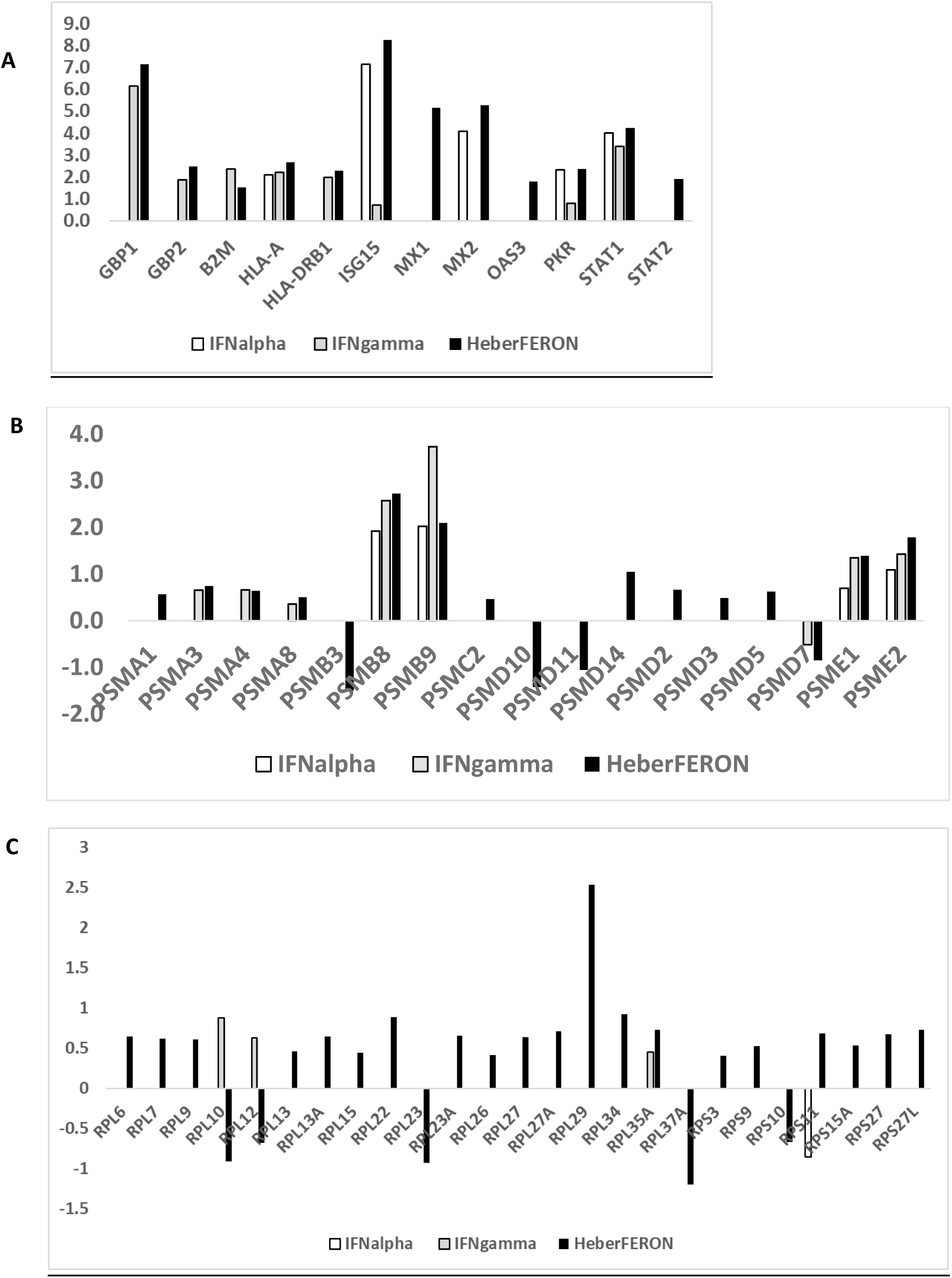
Changes in the expression of selected proteins regulated by IFNα (alpha) and IFNγ-(gamma) and HeberFERON. Graphs show the Fold change (FC) respect to untreated cell control for (A)- “Classical” proteins regulated by Interferons, (B)- Proteasome components, (C) Ribosomal proteins.

**Supplementary Fig. 2S.**
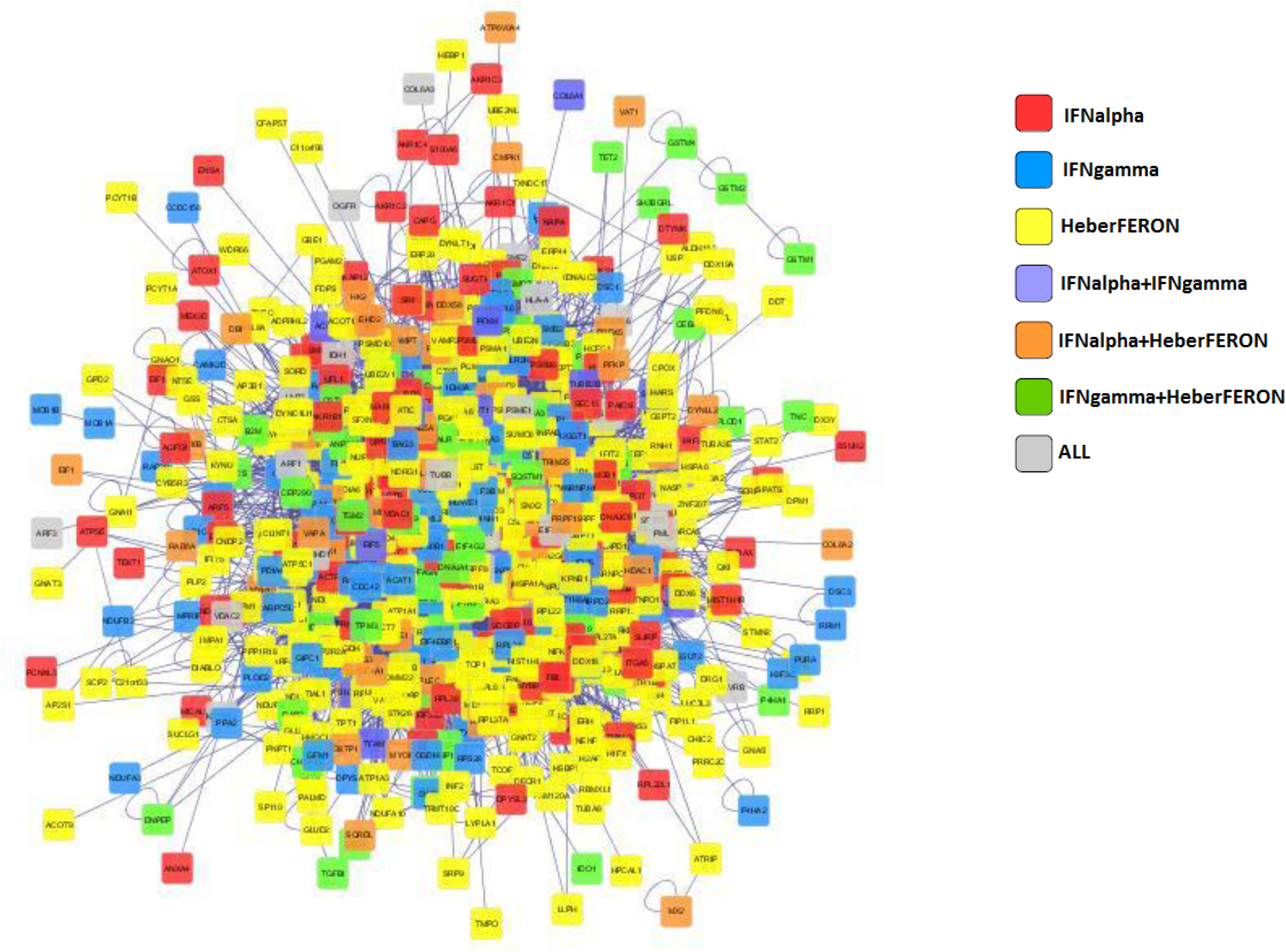
Network connection among ALL proteins differentially expressed in the experiment. Cytoscape framework was used to build the network using Bisogenet as a Plugin and colored nodes to explain treatment that regulated each protein in the network as indicated in the legend.

## Notes

### Competing Interest Statement

The authors have declared no competing interest.

